# REM: An Integrative Rule Extraction Methodology for Explainable Data Analysis in Healthcare

**DOI:** 10.1101/2021.01.22.427799

**Authors:** Zohreh Shams, Botty Dimanov, Sumaiyah Kola, Nikola Simidjievski, Helena Andres Terre, Paul Scherer, Urška Matjašec, Jean Abraham, Mateja Jamnik, Pietro Liò

## Abstract

Deep learning models are receiving increasing attention in clinical decision-making, however the lack of explainability impedes their deployment in day-to-day clinical practice. We propose REM, an explainable methodology for extracting rules from deep neural networks and combining them with rules from non-deep learning models. This allows integrating machine learning and reasoning for investigating basic and applied biological research questions. We evaluate the utility of REM in two case studies for the predictive tasks of classifying histological and immunohistochemical breast cancer subtypes from genotype and phenotype data. We demonstrate that REM efficiently extracts accurate, comprehensible rulesets from deep neural networks that can be readily integrated with rulesets obtained from tree-based approaches. REM provides explanation facilities for predictions and enables the clinicians to validate and calibrate the extracted rulesets with their domain knowledge. With these functionalities, REM caters for a novel and direct human-in-the-loop approach in clinical decision-making.

## 1 Introduction

Diagnosis, prognosis and treatment planning in healthcare are nowadays informed by a variety of data types ranging from imaging to genomics biomarkers and electronic health records. Machine learning (ML)^1^, and in particular deep neural networks (DNNs) are capable of handling such high volume of heterogeneous data. However, while accurate, the opacity of many ML models (e.g., DNNs) poses a great challenge for their deployment in safety critical domains, such as oncology.

Dealing with ML opacity issues has given rise to the emerging fields of interpretable and explainable ML, which have been the subject of much debate^1^ in the community in recent years^2^. Interpretable ML focuses on the model itself and “how” well its underlying processes for making a prediction can be understood. Explainable ML, on the other hand, solely focuses on understanding “why” an ML model makes a prediction. Explanation is readily available in interpretable models (e.g., decision trees, rulesets). For non-interpretable models (e.g., DNN, SVM), the explanation is often provided by post-hoc means^3–6^, most common of which is feature importance^7^, where for each prediction the importance of individual features is approximated^8–10.^ Recent work^11–13^, however, has shown that feature importance methods are fragile and susceptible to adversarial attacks. In addition, recent user studies have shown that feature importance does not necessarily increase human understanding of the model and its predictions^14,15^, making it a unideal candidate for clinical domains.

To address this problem we use a combination of ML and machine reasoning (MR). This combination enables explainable data analysis in a way that facilitates the involvement of human experts in the analysis and the clinical decision-making based on this analysis. In particular, we propose an integrative Rule Extraction Methodology (REM) (Figure 1), that via its main component (REM-D), approximates a DNN with an interpretable ruleset model and uses that ruleset to explain the predictions of DNN. For approximation, REM-D decomposes a DNN into adjacent layers from which rules are extracted and merged to map input to output. Compared to the methods that extract rules from the network predictions directly without considering the inner working of the network, decompositional approaches benefit from the noise removal property of neural networks and the hierarchical representations of data learnt by the network through the composition of nonlinear transformations from one layer to the next^16–18^.

**Figure 1.**
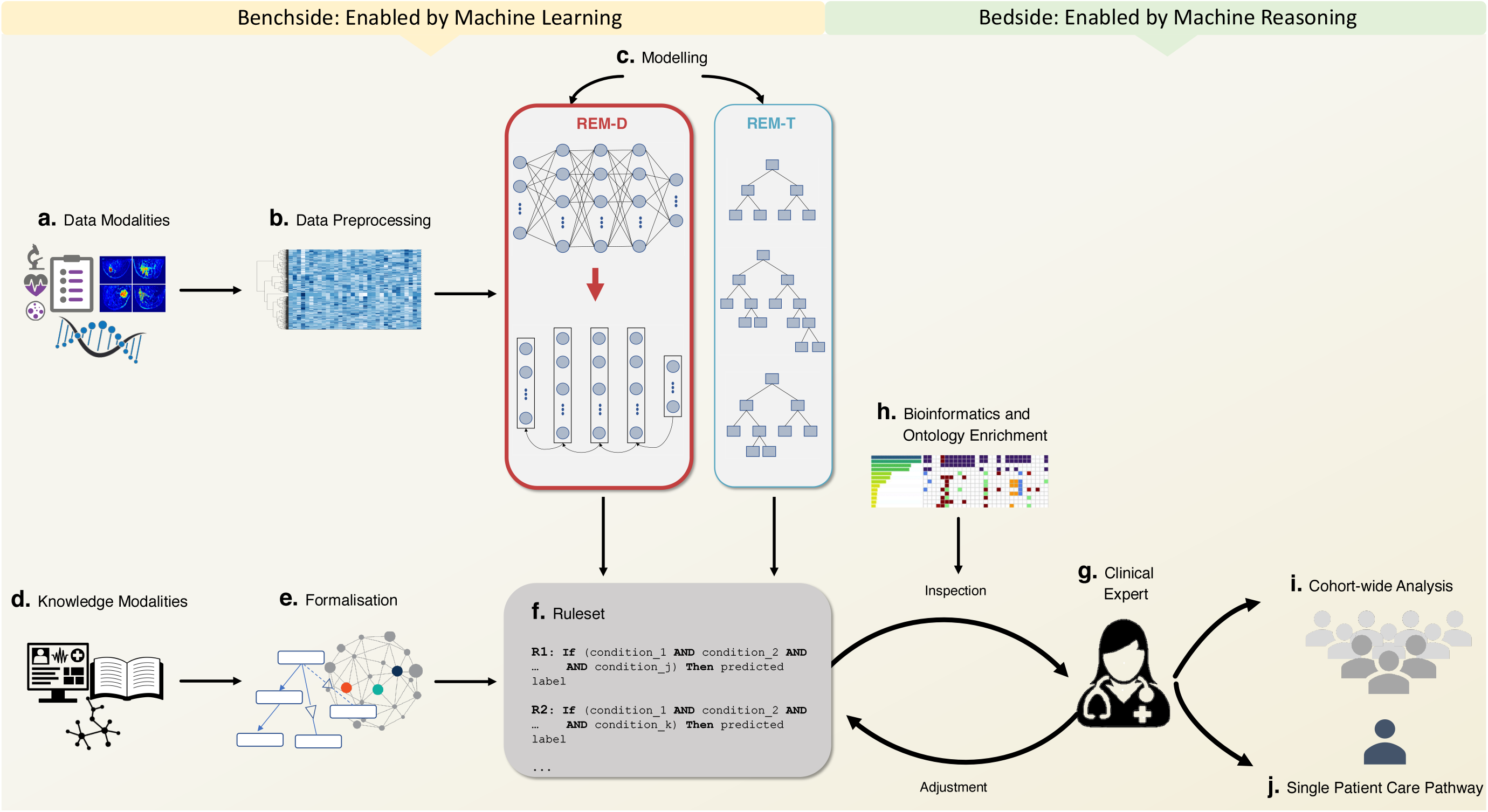
Workflow of REM for assisting bench-to-bedside research, a process in which research results are translated into new ways of treating patients. **a**. REM gets tabular input from various modalities (e.g., genomics, imaging, clinical). **b**. These modalities are subject to preprocessing and feature selection (e.g., using bioinformatics tools). **c**. REM-D and REM-T extract rules from trained DNNs and tree-based approaches used for modelling data modalities. **d**., **e**. Rules extracted from knowledge modalities (e.g., medical guidelines, electronic health records), and formalised (e.g., using knowledge graphs and ontologies) can be integrated into the rules extracted from data modalities. **f**. The final extracted ruleset allows reasoning to predict a target using multiple data, multiple knowledge, or a combination of data and knowledge modalities. **g**., **h**. Experts ^*a*^ can inspect the biological relevance of the ruleset using existing bioinformatics and ontology enrichment tools. They can also adjust it with their expertise and do further inspection to check the impact of such adjustments. **i**. Rulesets can be used by experts for analysis at the cohort-wide level to check population wide hypothesis and identify subgroups. **j**. Similarly, they can be used by experts for making a prediction for a single patient that impacts their care pathway, while accessing an explanation for the prediction.

The main motivation behind approximating DNNs with ruleset models, as opposed to other interpretable models, is in easy integration of such rulesets with other data and knowledge driven rules to allow multi-modality reasoning, a common practice in oncology. As an instantiation of combination with other data-driven rules, the REM-T component of REM extracts rulesets from tree-based approaches (e.g., decision trees, random forests) that can be combined with rulesets coming from the REM-D component, all of which can similarly be combined with rulesets from knowledge such as medical guidelines.

The final ruleset, as an end product, can easily be deployed in clinical decision support systems to enable clinicians to analyse the data and make predictions, and get explanations for the predictions. It also caters for verifiability and simulatability. With the former, clinicians can inspect the final ruleset model and its predictions to verify biological relevance in conjunction with bioinformatics tools. The latter reveals the impact that an input perturbation may have on the model’s prediction, thus allowing clinicians to adjust the model and its predictions based on their expertise.

Via two case studies that investigate a basic and an applied biological question using METABRIC dataset^19^, a cohort of 1980 breast cancer patients, we demonstrate the utility of REM in assisting bench-to-bedside oncology research, a process via which the results of basic biological research on data collected during trials is used to reinvent treatment pathway for patients. On the benchside, enabled by machine learning, the data and knowledge is integrated (Figure 1, a-f), and on the bedside, enabled by machine reasoning, the result of integration is made available to clinicians who can use it for population wide or single patient care (Figure 1, g-j).

The overall REM pipeline makes the following contributions to bench-to-bedside research: (i) it provides a flexible way to combine deep learning with non-deep learning models; (ii) it highlights the similarities and differences between the rules of decision-making by a machine and human experts; (iii) it caters for a novel and direct human-in-the-loop approach in clinical decision-making.

## 2 Results

We employ REM in two real world case studies on breast cancer for predicting imaging histological subtypes based on mRNA expression data and predicting immunohistochemical (IHC) subtypes based on a combination of clinical and mRNA expression data. The former task is focused on investigating a basic research question about the connection between genomics and imaging data, while the latter is focused on the applied question of IHC subtypes which is crucial in patient care pathway (e.g., whether they can benefit from hormone therapy as a part of their treatment pathway).

The results of rule extraction using the REM-D component are evaluated from three quantitative perspectives: (i) predictive performance (i.e., accuracy and fidelity of the ruleset), (ii) comprehensibility (i.e., size of ruleset and rules average length), and (iii) efficiency of rule extraction (i.e., time and memory usage). We demonstrate that in a time and memory efficient manner REM-D extracts ruleset models that are accurate and comprehensible. In addition to these quantitative measures, we show how the rulesets extracted by REM-D can be (i) inspected using bioinformatics tools, (ii) adjusted based on experts’ domain knowledge, and (iii) integrated with rules coming from other modalities and models.

### 2.1 Case studies

#### Case study I: Predicting histological subtypes of breast cancer from mRNA expressions

Histological subtypes of breast cancer aim to capture the heterogeneity of breast cancer based on the morphology evident in the pathology images of tumours. Invasive Ductal Carcinoma (IDC) and Invasive Lobular Carcinoma (ILC) are the two most common histological subtypes of breast cancer identified from pathology images, respectively. There is little known about the connection between genomics factors and these histological subtypes^20, 21^. To reveal the connection between the two, we use what we refer to as cross-modality reasoning: reasoning involved in predicting a target that is based on different data modalities rather than those inputted to the model. To this end, we predict the histological subtypes of IDC and ILC from mRNA expression profiles of patients in the METABRIC dataset^19^, using DNNs (Figure 2a). 1694 patients out of the total 1980 in METABRIC belong to one of these two subtypes. Using 80% of these 1694 records for training (1355 patients) and 20% (339 patients) for testing, the predictive performance of a neural network is measured across five folds, when identifying IDC vs ILC subtypes from 1004 mRNA expressions.

**Figure 2.**
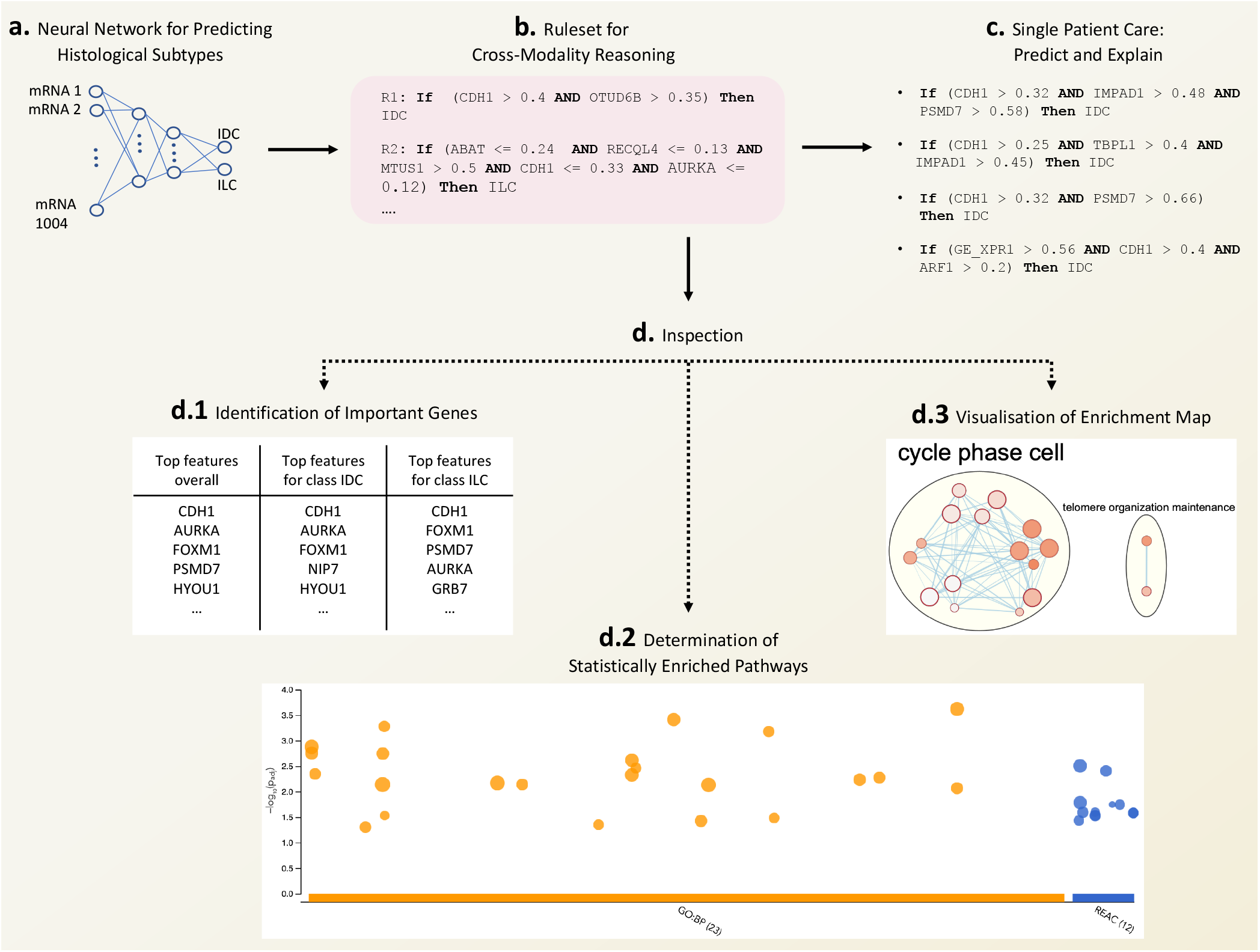
Case study I - an instantiation of REM pipeline (Figure 1) with a single data modality on the benchside, and “Inspection” functionality and “Single Patient Care” on the bedside. **a.** A neural network trained to predict histological subtypes of breast cancer based on 1004 mRNA expressions. **b.** Ruleset extracted from the trained network using REM-D to be used for single patient and cohort-wide analysis. **c.** The explanation for predicting the histological type of a hypothetical patient (generated by randomly sampling numbers between 0 and 1 to represent the input features) in the form of specific rules used for this prediction. **d.** Inspection of biological relevance of top recurring genes in the rulesets by ontology enrichment analysis. The details of pathways enriched (orange and blue circles) can be found in Supplementary Table 8.

#### Case study II: Predicting IHC subtypes of breast cancer from mRNA expressions

Apart from histological subtypes, patients in the METABRIC dataset are assigned to various other groups, one of which is the two IHC subtypes (ER+ and ER-) that are very important in deciding treatment options. Similar to previous scenario, using 80% of the METABRIC dataset for training (1584 patients) and 20% (396 patients) for testing, the predictive performance of a neural network is measured across five folds, when identifying patients IHC subtypes from 1000 mRNA expressions (Figure 3a).

**Figure 3.**
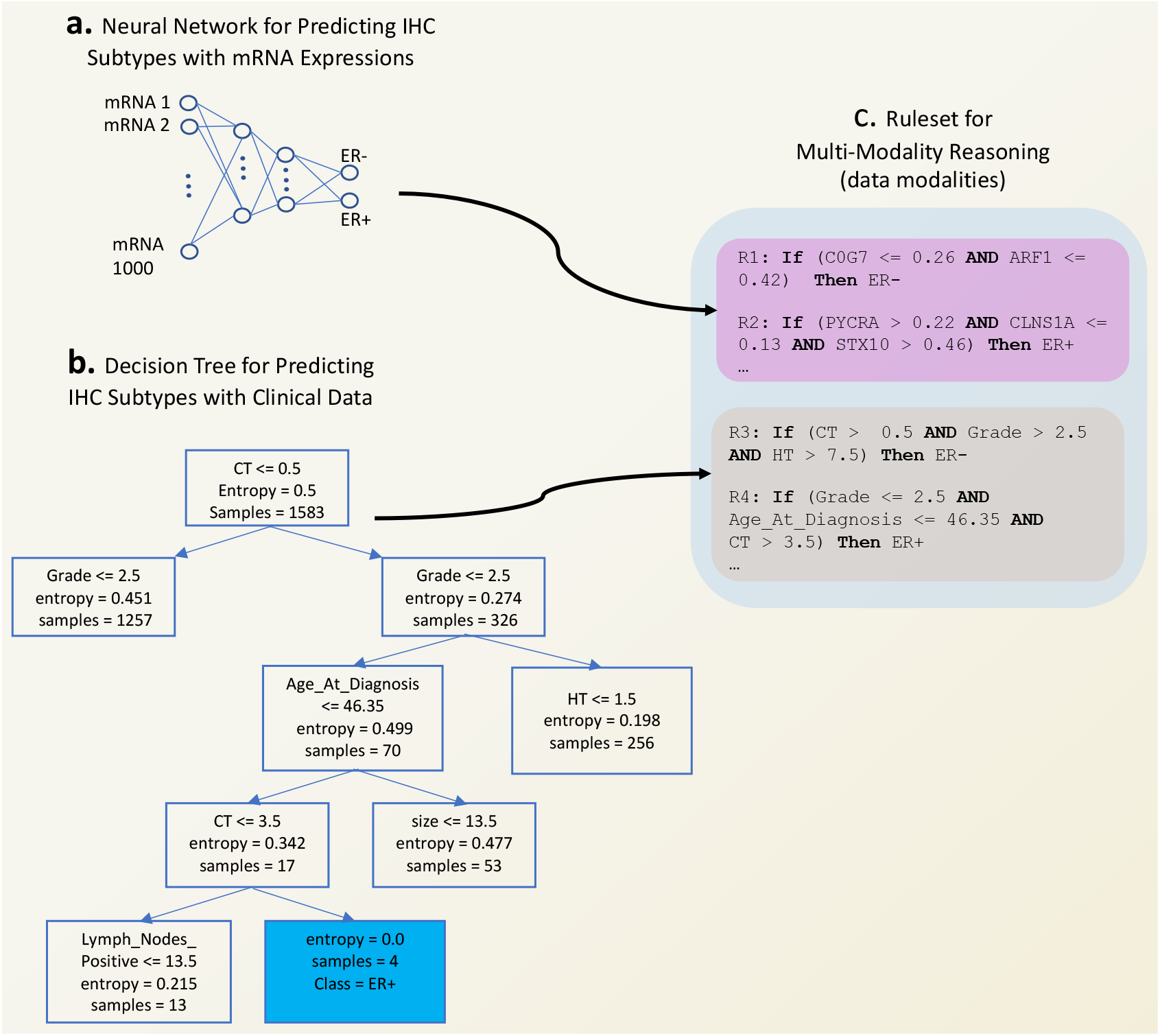
Case study II - an instantiation of REM pipeline (Figure 1) with two data modalities on the benchside. **a.** A neural network trained to predict IHC subtypes of breast cancer based on 1000 mRNA expressions. **b.** A decision tree trained to predict IHC subtypes of breast cancer based on 13 clinical variables. **c.** Combination of rulesets extracted from mRNA expressions using REM-D and from clinical variables using REM-T that gives rise to a ruleset for multi-modality reasoning to be used for single patient and cohort-wide analysis.

### 2.2 REM-D efficiently extracts accurate and comprehensible rulesets

Using REM-D, we extract rules from neural networks for both case studies above (Figure 2b and Figure 3c). The results in terms of predictive performance, efficiency and comprehensibility are reported in Table 1.

**Table 1.** Rule extraction from neural network when predicting ILC/IDC and ER+/ER-using mRNA expressions. Results reported are average across five fold cross validation for accuracy and fidelity, and median for the rest of the metrics. Results for each fold for ILC/IDC and ER+/ER- predictions are in Supplementary Tables 4 and 5, respectively.

| target | DNN accuracy | REM-D accuracy | REM-D fidelity | REM-D duration (sec) | REM-D memory (MB) | size of ruleset | rules average length |
| --- | --- | --- | --- | --- | --- | --- | --- |
| ILC/IDC | 0.882 | 0.886 | 0.884 | 143.792 | 573.231 | 137 | 4.3 |
| ER+/ER- | 0.957 | 0.921 | 0.926 | 100.714 | 349.951 | 103 | 5.1 |

For the first case study, from predictive performance perspective, the extracted rules closely mimic the decision of the neural networks (average fidelity of 88.4%). The average accuracy of the rules when used for prediction is 88.6%, which is almost identical to the original neural network. Comprehensibility remains high too, making the ruleset easy to audit and comprehensible for clinicians: the median of the size of the ruleset is 137, while the median of the average length of rules is 4.3 (i.e., rules have between 4 to 5 conditions). With respect to efficiency, rules are extracted efficiently in below 3 minutes (median) with memory consumption of 573.231 megabytes (median), using the hardware stated in Section 4.2. Note that the standard deviation across folds for comprehensibility and efficiency metrics can be high. Therefore, for these metrics we report the median instead of average, which gives a clearer overview of the REM-D behaviour. We outline our hypothesis about this observation in the Discussion section.

Regarding the second case study, the average accuracy and fidelity of the rules when used for prediction are 92.1% and 92.6%, respectively. This means that replacing the neural network with the ruleset compromises just over 3% of the accuracy. The median of the size of the ruleset is 103, while the median length of rules is 5.1, ensuring a high degree of comprehensibility. The efficiency remains high. It takes below 350 megabytes and 2 minutes to extract the rules, using the same hardware.

Given the high fidelity and accuracy of the extracted rulesets, when acting as a surrogate to the original model, they can be used for both cohort-wide analysis and single patient care planning (Figure 1i, j). On the cohort level, use cases include analysing the overall ruleset to give an overview of features most frequently used in rules for a whole cohort or subgroups within a cohort (e.g., IDC vs ILC patients or ER+ vs ER-patients in case studies I and II). Similarly, the threshold of features used in rules can reveal information about the cohort (e.g., whether a gene expression value is consistently below or above a level for a subgroup versus another - although the absolute gene expression values as they appears in the rules may not be readily biologically meaningful, analysing their overall patterns may point towards their consistent over- or under-expression for a certain condition, a very meaningful biological theme).

On the single patient level, rulesets can be simulated easily to get a prediction for a new patient. In addition, it can be explained why a prediction was made, where the explanation is in the form of rules that were satisfied (i.e., the conditions in them were met) and thus counted towards the prediction (e.g., Figure 2c). Clinicians can also potentially perturb the patients features (e.g., decrease their BMI) to check the impact of such change on the prediction of the model (e.g., checking whether encouraging the patient to decrease their BMI changes their treatment outcome for the better).

Having assessed the extracted ruleset from quantitative perspective, in the next three sections, we focus on the qualitative aspects. Each aspect is contextualised for one of the case studies and can be applied to the other case study in a similar fashion.

### 2.3 Extracted rulesets can be inspected using bioinformatics tools

Depending on the data modality, inspecting and verifying biological relevance of the rules may be challenging. This is in particular the case when working with genomics data that is hard to interpret. Here we show how the interpretation can be done by relying on bioinformatics tools. To inspect the biological relevance of the rules extracted for the case study I, we follow the protocol in Reimand et al.^22^ that is based on well-established bioinformatics tools designed to interpret gene lists resulting from experiments. Figure 2, parts d.2, and d.3 show the two major steps of the protocol after defining the gene list of interest in d.1. These steps are: determining statistically enriched pathways (d.2), and visualising and interpreting the results (d.3). In line with the protocol recommendation we use g:Profiler^23^ and EnrichmentMap^24^ for identifying pathways enriched in the gene list of interest relative to what is expected by chance, and mapping them visually to facilitate identifying the main biological themes they represent. As a proof of principle, we query the top 50 genes that appear in the rules for class IDC in g:Profiler for the pathways they enrich in biological processes of GO^25^ and molecular pathways of Reactome^26^. The results are then passed on to EnrichmentMap for visualisation and annotation. Supplementary Table 8 and Figures 7 and 8 include details on steps d.1, d.2 and d.3, respectively. A systematic approach to integrating rule extraction with bioinformatics, such as the protocol followed, gives clinicians a biological insight to what the rules have captured and how biologically relevant that is to the task in hand.

In addition to genes, the rules include information about their expression, which may point toward the association of a phenotype with over- or under-expression of some genes. In the case reviewed above, CDH1 is the most common feature. This gene codes for a cadherin involved in the key Wnt-mediated beta catenin signaling. Close to 80% of the rules associated with class ILC that include this feature prescribe an upper bound restriction for it, which is aligned with the reported association of ILC tumour and the under-expression of CDH1^20, 21^. In contrast, all the rules associated with class IDC that include this feature restrict its lower bound. This means that these rules are more likely to get triggered and therefore predict IDC for patients with over-expression of CDH1. In addition to CDH1, the three features that are commonly used in rules for both classes are AURKA, FOXM1, HYOU1. All rules for class IDC restrict the lower bound for AURKA, FOXM1 and HYOU1. This is likely to point towards the over-expression of these genes in IDC tumours. The opposite is true for rules associated with class ILC: close to 99% and 95% of the rules limit the upper bound of AURKA and FOXM1, respectively; as well as all the rules limiting the upper bound of HYOU1, making a case for potential under-expression of FOXM1 and HYOU1 in ILC tumours. The connection between the boundaries of gene expression in the rules and various phenotypes can be explored in the same fashion for the rest of the genes.

### 2.4 Extracted rulesets can be adjusted based on domain knowledge

There are various ways of incorporating domain knowledge expressed by clinicians. One way is for clinicians to express important features to focus on. This information can be used to impose a hierarchy on the ruleset by assigning scores to rules, where the score takes into account the inclusion of the preferred expressed features. We use an augmented hill climbing algorithm for scoring rules. In the base version^27^ the algorithm assigns a score to each rule based on its coverage, accuracy and length, where high accuracy and coverage are rewarded, while high score for length is penalised. In the augmented algorithm, referred to as personalised ranking^28^, the rules that include features proposed by clinicians get additional scores and are thus likely to rank higher in the hierarchy of rules (Equation 2, Methods). To show the impact of scoring, we pick four genes at random (e.g., KCTD3, RARA, STARD3 and ERLIN2) and assume they are expressed as highly relevant by an expert when predicting IHC subtypes using 1000 mRNA expressions (Figure 3a). Note that we used random selection to show the generality of the ranking. In a real world setting the random genes can be replaced by, for instance, known cancer driver genes. The frequency of favourite features in the top 70% of the ruleset increases when using personalised ranking (purple bars, Figure 4) as opposed to the ranking merely based on coverage, accuracy and length (blue bars, Figure 4), indicating that the model is indeed adjusted in the direction of expert knowledge. ERLIN2 is the exception, where the personalised ranking does not make a difference. Potential reasons for this are the absence of more rules that include ERLIN2. High specificity of the rules that include ERLIN2 could be another reason: although the ranking favours the inclusion of this feature, it does not favour long rules that are highly specific.

**Figure 4.**
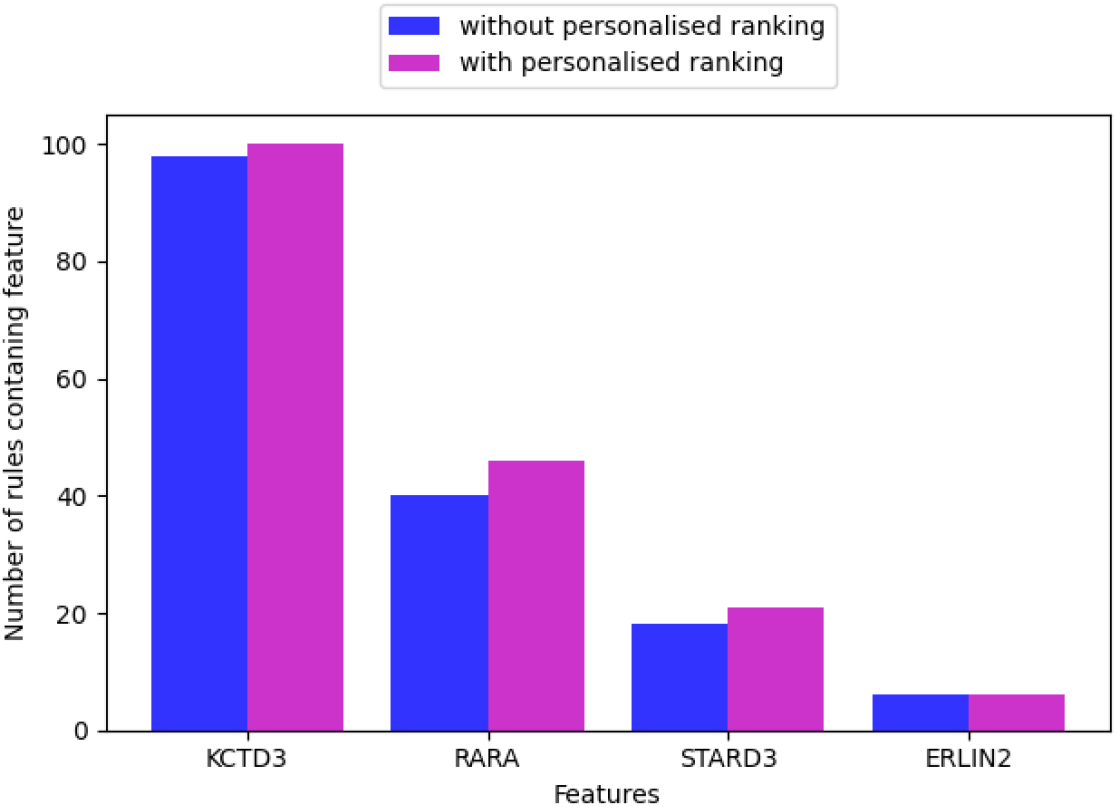
An instantiation of “Adjustment” functionality within REM pipeline (Figure 1): The impact of personalised ranking on the increase in the number of higher rank rules (top 70% of the rules) including the preferred features expressed by clinicians.

### 2.5 Extracted rulesets can be integrated with rulesets from other modalities and models

Beside mRNA expressions, IHC subtypes can also be predicted from clinical data in METABRIC. Unlike the genomics part of the data that tends to be high dimensional, the clinical part often consists of a handful of variables. We predict the IHC subtypes from clinical data using a decision tree (Figure 3b). The average accuracy of decision trees across the folds is 81%, while the median of the ruleset size and rule length are 29 and 4.6, respectively, as presented in Table 2, second row.

**Table 2.** Integration of rules extracted from mRNA data modelled using a neural network (mode REM-D) with rules from clinical data modelled using a decision tree (mode REM-T), when predicting IHC subtypes. Results reported are median across five fold cross validation. Results for each fold are in Supplementary Tables 5, 6 and 7 for mRNA, Clinical, mRNA + Clinical, respectively.

| data modality | rule extraction mode | rule extraction accuracy | size of ruleset | rules average length |
| --- | --- | --- | --- | --- |
| mRNA | REM-D | 0.921 | 103 | 5.1 |
| Clinical | REM-T | 0.81 | 29 | 4.6 |
| mRNA + Clinical | REM-D + REM-T | 0.933 | 133 | 5.0 |

To allow multi-modality reasoning, we combine the rules extracted from the decision tree, using REM-T, with the rules from the neural network, using REM-D (Figure 3c). As displayed in Table 2, third row, this increases the accuracy of the ruleset slightly over the five folds, in comparison with the ruleset that relies on rules extracted from the neural network only. The size of the ruleset increases as expected, while the median of the average length of the rules drops below that of the rules extracted by REM-D. More important than the slight increase in the accuracy is the fact that rules from different modalities modelled with various models can indeed be integrated and contribute to one another.

## 3 Discussion

Heterogeneity and high volume of data in domains such as oncology challenge common statistical methods. Although DNNs has shown promise in overcoming these challenges, they come with their own challenges due to lack of explainability. This is evidenced by the scepticism of clinical community about ML systems^29,30^ as well as the growth of interpretability and explainability methods used in healthcare to address this problem. The majority of these methods are local, pointing to important input features that were most influential in a prediction^7,31–33^. Apart from vulnerability to adversarial attacks^11–13^ and scalability issues caused by the need for model retraining^8^ to generate explanation, user studies do not support feature importance explanatory utility for human users^14, 15^. REM, the pipeline proposed here for explainable data analysis, provides a global view of explainability by combining ML and MR. The ruleset models extracted by REM give an overview of the original underlying model, while remaining usable for local explanation of individual predictions.

In addition to providing explainability, the rule-based nature of REM makes it a unique candidate for use in healthcare, due to facilitating integration of heterogeneous data and incorporation of human-in-the-loop. Instead of using a single model for all modalities, each modality is modelled using the prediction model that suits the data best, while the integration happens at the rule level to allow multi-modality reasoning. As a result, deep learning and non-deep learning approaches can flexibly be used in combination; and uninterpretable models such as DNNs are only used when they outperform more interpretable models such as decision trees (e.g. See Supplementary Table 3) and are avoided otherwise. Once the rules are extracted and integrated, clinicians can contrast them with their domain knowledge, manipulate them, and also impose hierarchy on them to emphasise the subset of rules that may be more useful for a subgroup of patients (e.g., of specific ethnicity).

**Table 3.** A performance comparison between Decision Tree, Random Forest and Neural Network with 1004 mRNA expressions as input and ILC/IDC as output.

| fold | decision tree |  | random forest |  | neural network |  |
| --- | --- | --- | --- | --- | --- | --- |
|  | accuracy | AUC | accuracy | AUC | accuracy | AUC |
| 0 | 0.897 | 0.718 | 0.918 | 0.639 | 0.903 | 0.811 |
| 1 | 0.879 | 0.723 | 0.918 | 0.609 | 0.906 | 0.813 |
| 2 | 0.846 | 0.666 | 0.929 | 0.617 | 0.870 | 0.804 |
| 3 | 0.876 | 0.698 | 0.905 | 0.589 | 0.908 | 0.731 |
| 4 | 0.885 | 0.624 | 0.914 | 0.578 | 0.825 | 0.78 |
| average | **0.877** | **0.686** | **0.917** | **0.606** | **0.882** | **0.788** |

We demonstrated the use of REM and its functionalities in two breast cancer case studies. In the case study I (Figure 2) we used genomics data to predict image-based targets. In the case study II (Figure 3) we looked at the complementarity of modalities (e.g., mRNA expression and clinical) for predicting certain targets (e.g., IHC subtypes), and showed that modalities can indeed contribute to one another when making predictions. Similar to these case studies, rules can be extracted from knowledge modalities (e.g., electronic health records and biomedical ontologies) and integrated with data-driven rules. Figure 5 showcases such a scenario. Rules extracted from data provide input to rules extracted from the Breast Cancer Staging Ontology^34^, which is based on the latest AJCC Cancer Staging Manual^35^. In a hypothetical scenario, where the IHC subtypes and tumour stage of the patients are unknown, the former can be predicted from data modalities (as demonstrated in case study II, Figure 3d) and used in the prediction of the latter, assuming other variables required for this latter prediction (i.e., Grade, HER2 status, PR status, T: severity of tumour size, N: severity of the spread to the lymph nodes, M: metastasize status) are known and provided as facts.

**Figure 5.**
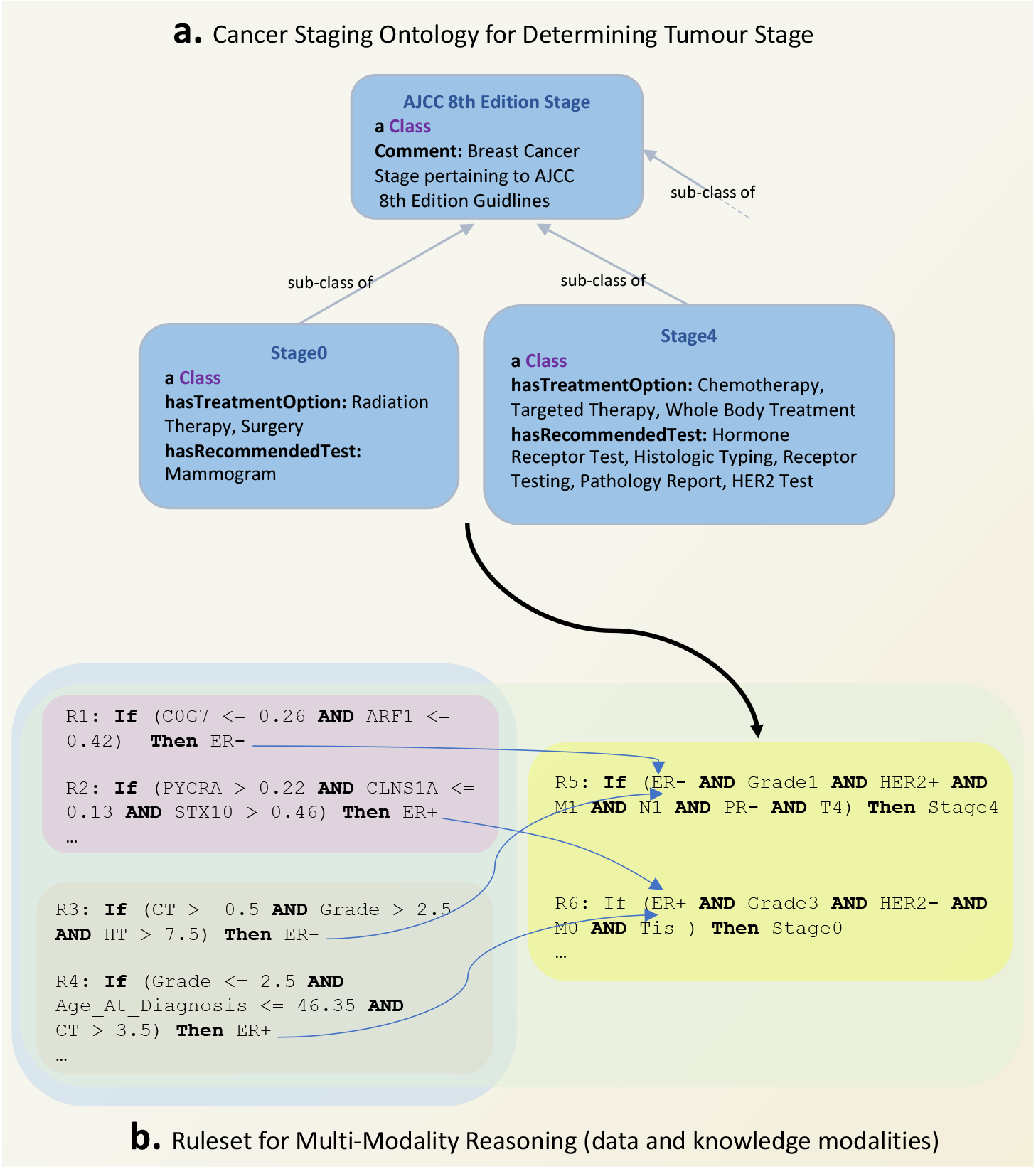
An instantiation of REM pipeline (Figure 1) with a combination of data and knowledge modalities on the benchside. **a**. An existing ontology that formalises the medical guidelines in the latest AJCC Cancer Staging Manual. **e**. Rulesets from ontological knowledge (yellow box) integrated with rules from data modalities (pink and grey boxes).

Rule extraction for interpretability purposes is very common in tree-based approaches^36–38^, however rule extraction from neural networks is rare, with the majority of methods being limited to rule extraction from networks with one hidden layer^18^. Zilke et al.^39^ extract rules from networks with more than one hidden layer (i.e., DNNs), however, the approach has scalability issues due to prohibitively high memory and time consumption^4, 39^. Due to these reasons, it is not possible to directly compare Zilke et al.^39^ to REM-D, however, we discuss the algorithmic differences between the two that are crucial in time and memory consumption in the Methods section. Another alternative for rule extraction from DNNs is that proposed by Nguyen et al.^40^. This approach is restricted to networks that use ReLU as the activation function between hidden layers, and is therefore not generally applicable. Kazhdan et al.^41^ propose another decompositional approach to extract information from Convolutional Neural Networks (CNNs) layers in the form of “concepts”. Concepts, in the setting of images, are high-level semantic units (e.g., lung opacity, foreign object) rather than individual input features (e.g., pixels, characters). While, using REM-D to extract rules from image-based data modalities that are based on pixel-level features may not prove informative, image patches or concepts can provide a great medium for REM-D to extract rulesets from CNNs that are based on high level units, meaningful to humans^11^. In the case of other architectures such as recurrent neural networks (RNNs), similar to^42^, REM-D can extract rulesets at every time step followed by a merge step to provide a global view of the underlying model. However, in contrast to^42^, which provides transition rules that only describes the relationship between the output and the last hidden layer, REM-D provides a hierarchy of rules across the entire architecture to provide explanations about the relationship between the inputs, hidden layers, and outputs.

We envision several exciting avenues for future development of REM on theory, application and biological relevance fronts. Regarding the technical side, a limitation is when for large models the rule extraction is intractable or the number of rules extracted is so high that it is difficult to claim interpretability. We demonstrate the complexity of rule extraction in the Methods section and discuss ways of improving it. Related to number of rules, deep learning models often have multiple optima of similar predictive accuracy, while their interpretability can vary considerably^43^. We postulate that neural networks that give rise to a smaller number of rules in a shorter span of time and with less memory consumption, when subjected to rule extraction, are those that are more interpretable and hence easier to explain with a smaller number of symbolic rules. Our results (Supplementary Table 4 and Table 5) also suggest that the larger rulesets tend to have longer rules that are highly specific due to inclusion of several conditions and are therefore less suited to generalise to unseen samples. To address this limitation and control the number of rules extracted, we plan to use optimisation techniques that focus on optimising deep models for human-simulatability^44^. Such optimisation is expected to encourage finding models whose decision boundaries are well-approximated by small decision trees that in turn give rise to a small ruleset, which are less likely to overfit. Related to facilitating the presentation of rulesets is providing visualization mediums. One way to realise this goal is to use existing algorithms that learn trees from rulesets^45^, instead of directly from training data, to map rulesets to diagnostic decision trees that are easy to visualise.

**Table 4.** Rule extraction from neural network with 1004 mRNA expressions as input and ILC/IDC as output.

| fold | DNN<br>accuracy | REM-D<br>accuracy | REM-D<br>fidelity | REM-D<br>duration (sec) | REM-D<br>memory (MB) | size of<br>ruleset | rules<br>average length |
| --- | --- | --- | --- | --- | --- | --- | --- |
| 0 | 0.903 | 0.888 | 0.903 | 143.792 | 573.231 | 155 | 4.3 |
| 1 | 0.906 | 0.885 | 0.891 | 32.06 | 146.538 | 6 | 1.8 |
| 2 | 0.870 | 0.867 | 0.861 | 143.881 | 642.576 | 137 | 7.7 |
| 3 | 0.908 | 0.902 | 0.929 | 59.239 | 227.721 | 18 | 2.4 |
| 4 | 0.825 | 0.888 | 0.837 | 157.059 | 630.239 | 240 | 6.6 |
| average | **0.882** | **0.886** | **0.884** | **107.206** | **444.061** | **111.2** | **4.6** |

**Table 5.**
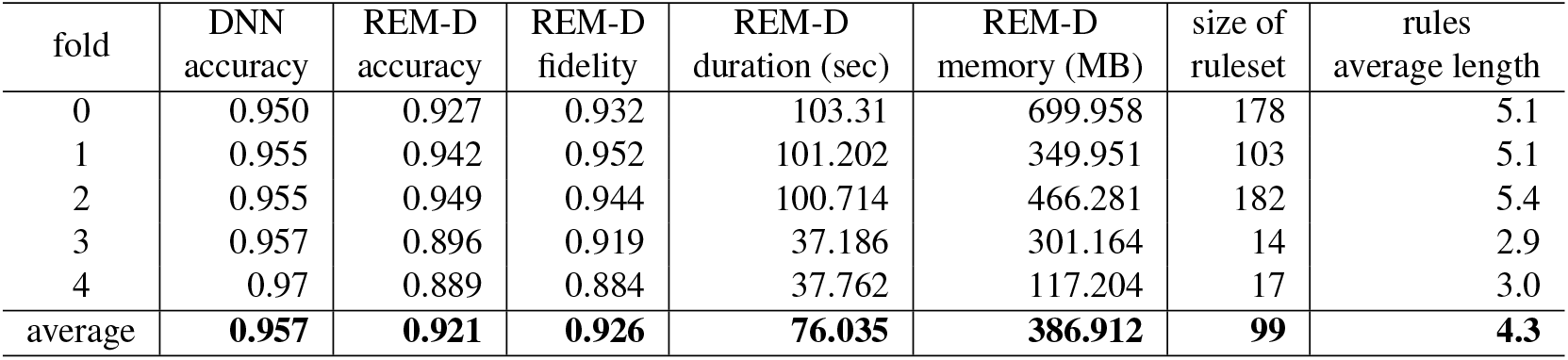
Rule extraction from neural network with 1000 mRNA expressions as input and ER+/ER-as output.

| fold | DNN<br>accuracy | REM-D<br>accuracy | REM-D<br>fidelity | REM-D<br>duration (sec) | REM-D<br>memory (MB) | size of<br>ruleset | rules<br>average length |
| --- | --- | --- | --- | --- | --- | --- | --- |
| 0 | 0.950 | 0.927 | 0.932 | 103.31 | 699.958 | 178 | 5.1 |
| 1 | 0.955 | 0.942 | 0.952 | 101.202 | 349.951 | 103 | 5.1 |
| 2 | 0.955 | 0.949 | 0.944 | 100.714 | 466.281 | 182 | 5.4 |
| 3 | 0.957 | 0.896 | 0.919 | 37.186 | 301.164 | 14 | 2.9 |
| 4 | 0.97 | 0.889 | 0.884 | 37.762 | 117.204 | 17 | 3.0 |
| average | **0.957** | **0.921** | **0.926** | **76.035** | **386.912** | **99** | **4.3** |

Application-wise, we believe that the deployment of REM for revealing the connections between various modalities can prove very useful when cost consideration makes accessing some data modalities (e.g., genomics) challenging. For example, genomics components involved in targeted gene panel testing, which are crucial in treatment planning, may be predicted based on widely available imaging data^46^. Investigation of use of REM for extracting rules of going from widely available data modalities (e.g., pathology imaging) to hard to access ones can be a step towards bringing personalised medicine to more patients. Finally, in terms of biological relevance, investigation of connection between proteins coded by genes in individual rules and sets of proteins participating in known pathways can shed light on the higher level biological functions that rules capture. We are positive that accommodating such functionalities creates more synergy for REM to be used in conjunction with other bioinformatics tools.

## 4 Methods

### 4.1 Data, pre-processing and feature selection

For all experiments we use a public breast cancer dataset of 1980 patients, METABRIC^19^, that aims to characterise breast cancer subtypes based on genomics and imaging data as well as clinical data.

#### Feature selection in case study I

The aim of this case study is to predict the two main histological subtypes of breast cancer, IDC and ILC (1547 IDC vs 147 ILC cases), using mRNA expressions. Feature selection from METABRIC for this task is done based on existing bioinformatics findings: we use the 1,000 putative breast cancer genes identified in Curtis et al.^19^, the significance of which is validated by revealing novel subgroups that have distinct clinical outcomes. These genes are identified based on a landscape created by integration of copy number aberrations (CNA) and expressions that highlights genomics regions which are likely to contain driver genes. Further bioinformatics findings show that in addition to distinct morphology, mRNA expression profiling of the two main histological subtypes demonstrates distinct molecular differences^20^. The main differences include the variant in expression of four genes: CDH1, MKI67, FOXA1 and PTEN. We add the mRNA expression of these four genes to the 1,000 putative ones (Figure 2a).

#### Feature selection in case study II

The aim of this case study is to predict the two main IHC subtypes of breast cancer, ER+ and ER-(1506 ER+ vs 474 ER-cases), using mRNA expressions and clinical data. Similar to the above case study, we use the mRNA expressions of the 1,000 putative genes (Figure 5a). The clinical data (Figure 5b) used in combination with the genomics data consists of 13 variables (age at diagnosis, tumour laterality, Nottingham Prognostic Index, menopause status, number of positive lymph nodes, chemotherapy agent, hormone therapy agent, radiotherapy agent, grade, tumour size, histological type, stage, cellularity).

There are no missing values in mRNA expressions. Expressions are normalised and scaled to [0,1] prior to use. Occasional missing values in clinical part are replaced by mean value for continuous variables. They are treated as a new category for categorical variables to represent an unknown value. After pre-processing, the data is sampled into five-fold cross validation splits based on the class distribution of the target.

### 4.2 Model selection

A fully connected neural network with two hidden layers is set up for the classification tasks of predicting histological subtypes and IHC subtypes outlined above. We used Keras library^47^ for this. Tanh is used as the activation function for the hidden layers, and softmax is used for the output layer. Adam optimiser^48^ is used for training without any regularisation. The weights of classes are considered too when fitting the model. The batch size, number of epochs, and the number of neurons in the two hidden layers are determined based on a grid search over the following parameters: batch-size ={16, 32, 64, 128}, epochs = {50, 100, 150, 200}, layer-1 = {16, 32, 64, 128} and layer-2 = {8, 16, 32, 64}. Best performance (i.e., average AUC of five-fold cross validation) is obtained with: batch-size=16, epochs=50, layer-1=128, and layer-2=16 when predicting histological subtypes using 1004 mRNA expressions. Incidentally, the same applies to predicting IHC subtypes. When modelling clinical data using a decision tree (Keras library^47^), rules were extracted without limiting the depth of the tree, as well as limiting trees to depths 5, 10 and 15 (max_depth= {5, 10, 15}). The splitting criterion used was “entropy”. The accuracy of predicting IHC subtypes was highest when maximum depth of the decision tree was limited to 5. All the experiments are done on a server with 66 GB RAM and 2 AMD 6367 processors.

### 4.3 REM-D rule extraction methodology

REM-D extracts rules (Algorithm 1) for a classification problem with *n* data points, 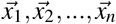, each with an associated class, *y*_*i*_ ∈ {*y*_1_, *y*_2_, …, *y*_*u*_}. Rules are extracted from a trained network with *k* hidden layers, {*h*_0_, *h*_1_, …, *h*_*k*_, *h*_*k*+1_}, where *h*_0_ and *h*_*k*+1_ are the input and output layers. Each layer *h*_*i*_ has *H*_*i*_ neurons. The input layer thus has as many neurons as the features available for each data point: 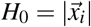. The number of neurons in the output layer equals to the number of classes: *H*_*k*+1_ = *u*. The activation values sampled at layer *h*_*i*_ for data point 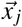 are denoted as 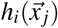.

#### Ruleset structure

Prior to outlining the methodology for rule extraction, we present the structure of the ruleset extracted by REM-D:

- Total ruleset: 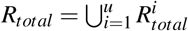
- Ruleset for each class: 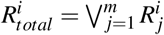
- Individual rules: 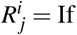 *antecedent* then *i*
- Antecedent: 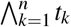
- Terms: *t*_*k*_: *h <*= *threshold* or *t*_*k*_: *h > threshold*

The total ruleset is the union of rulesets for each class. The ruleset for each class is a disjunction of individual rules, each of which is a conditional statement with an antecedent and the class they belong to as conclusion. The antecedent itself is a conjunction of conditional statements referred to as terms that are conditions on the activation value of neurons.

#### Rule extraction algorithm

In order to extract rules of the form described above from the network, REM-D first decomposes the trained network into adjacent layers and then uses C5.0^49^ to extract rules from pairs of layers in the network. Unlike CART^50^, in addition to tree induction, C5.0 is also a rule extraction algorithm^51^. Compared with other tree induction and rule extraction algorithms, C5.0 is faster than its predecessors C4.5^49^ - which itself is the successor to ID3^52^ - while consuming less memory. It also builds smaller rulesets that are more accurate. A comparison of C4.5 and C.50 by the author of both algorithms can be found in^53^.

When generating decision trees between layers, in order to identify the features that lead to a split, C5.0 uses entropy^54^ for measuring purity. The information gain of a feature is calculated based on the difference of entropy in the segment before the split and the partitions resulting from the split. The features with higher information gain are then used for splitting the data in the decision tree. Once the tree is generated, nodes and branches that have little effect on the classification errors are pruned. Rules are then extracted from the pruned tree and pruned further in accordance with their contribution to accuracy. The rules extracted by C5.0, from each two adjacent layers, *j* and *j* + 1, use the features from layer *j*. Thus, except when *j* is the input layer, the rules use features in a hidden layer. To make sure that the final ruleset maps the input to the output, rules are merged in a backward fashion, such that the hidden features in each rule are replaced by features in the previous layer until the input layer is reached and all rules are expressed only in terms of input features. Algorithmic details are as follows.

##### Algorithm 1 REM-D

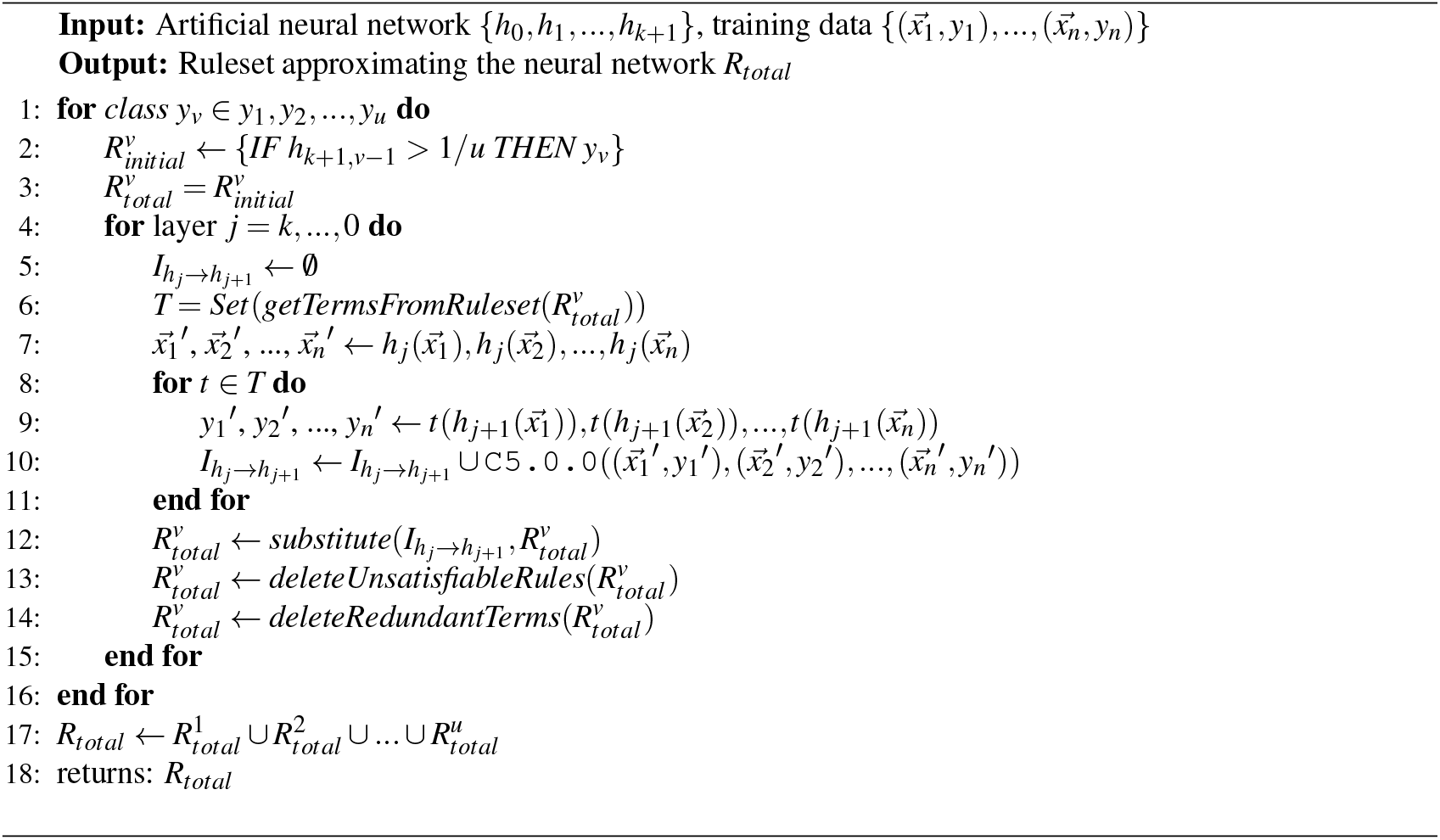

The algorithm starts by iterating over each class (line 1). For each class, a rule is defined that maps the output layer to the output classes (line 2). For example, if there are two classes, represented by neuron 0 and 1 in the output layer, which we assume is layer 5, for the first one we have: 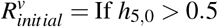 Then *y*_*v*_, where *h*_5,0_ refers to neuron 0 of layer 5. A total rule is introduced for each class that only contains the initial rule to begin with (line 3). For the class in hand, each layer preceding the output layer is considered in a descending order (line 4). For each layer and its proceeding layer an intermediate empty ruleset is formed that is going to be populated with rules extracted from these two layers (line 5). Unique conditions (referred to as Terms) that appear in the individual rules within the total ruleset are collected in a set (note that the set implementation guarantees the uniqueness of these conditions, and avoids the need for looking for redundancies and deleting them after the terms are collected) (line 6). These conditions are based on hidden neurons in the proceeding layer and need to be replaced with conditions based on the neurons in the current layer till we reach the input layer. For this first the activation values sampled from the current layer are noted (line 7). Next, each collected condition (line 6) is applied to the activations sampled from the proceeding layer to give a target value (line 9). Then in line 10, the values noted in line 7 and targets from line 9 are passed on to C5.0 algorithm for rule extraction. In the same line, the extracted rules will be added to the intermediate ruleset initiated in line 5. By substituting (Algorithm 2) the updated intermediate rules into the total class ruleset, the rules for the class are now described based on neurons in a layer one step closer to the input layer (line 12). Merging rules extracted from different layers may give rise to unsatisfiable rules that have contradictory conditions (e.g., Age_At_Diagnosis > 65, Age_At_Diagnosis <= 46) or rules with redundant conditions (e.g., Age_At_Diagnosis > 65, Age_At_Diagnosis > 67). Unsatisfiable rules are deleted in line 13, followed by deleting redundant conditions in line 14. For optimisation purposes, substitution, satisfiability and redundancy checking is done after each step of merging. This results in a lower number of shorter rules for the subsequent merge step, thereby improving time and memory usage. The procedure is repeated until the input layer is reached and the rules for each class are based on input features instead of neurons in the hidden layers. Finally, the rules that describe the behaviour of the network for each class in terms of input features are combined to give the overall ruleset (line 17) that is returned as output (line 18).

In order to make a prediction for a data point using the final ruleset, the majority vote is used: the ruleset for each class has a vote for the prediction which is essentially the number of rules within the class ruleset that are satisfied by the data point. The prediction for the data point is the class that has the majority vote (highest number of rules satisfied).

#### C5.0 parameters

The default parameter values are used in C5.0 (Algorithm 1, line 10). “winnowing” attribute is set to “True” and the number of “minCases” per leaf is determined by grid search. Winnowing in C5.0 works by calculating a feature importance for each feature based on error rate increase in the training set if the feature was excluded. When set to “True”, winnowing allows C5.0 to use only the important features for tree induction and rule extraction. The minCases parameter stops the decision tree from splitting further if the number of samples in each node drops below the set minimum number of cases after a split. For each experiment we extracted rules by setting the minCases values to: minCases= {5, 10, 15, 20} and chose the value that gave the most accurate final ruleset. When predicting histological subtypes, this value was 10, while 5 was the best value when predicting IHC subtypes.

##### Algorithm 2 Substitution Procedure

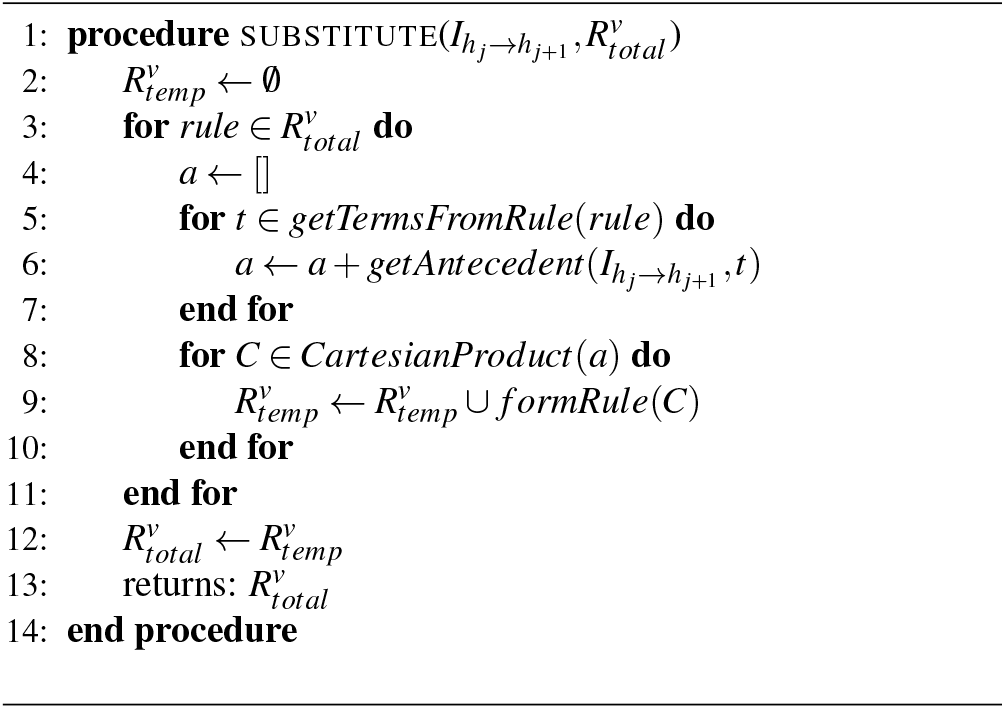

#### Substitution algorithm

The substitution procedure replaces the terms in individual rules of a class ruleset with the antecedent of the individual rules within the intermediate ruleset that has these terms as conclusion. For example, if we have *t* AND *t*′ → *y* and within intermediate ruleset we have the following two rules *a*→ *t* and *a*′→ *t*′, then the substitution gives rule *a* AND *a*′ *→ y*. This substitution essentially replaces the terms appearing in the rules of total ruleset with terms from a previous layer. As a result, the rules for the class are now described based on neurons in a layer one step closer to input layer.

The procedure starts with initiating a temporary ruleset for the class in hand (line 2). It then iterates over individual rules in the class total ruleset (line 3). For each of them an empty list is initiated (line 4). For each term in the rule (line 5), the intermediate ruleset is searched for rules with this specific term as conclusion. Those found are added to the list initiated earlier (line 6). If a rule has two terms *t*_1_ and *t*_2_, there may be several intermediate rules that have each term as conclusion, thus the list may look like 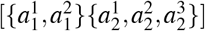, where 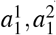 are antecedents for *t*_1_ and 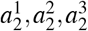 are antecedents for *t*_2_. Cartesian product of antecedent sets gives all possible combinations when substituting terms in rules within total ruleset (line 8). Each combination forms a new individual rule that is added to the temporary ruleset for the class (line 9). Once this procedure is repeated for each rule, the temporary ruleset replaces the total ruleset (line 12). The updated total ruleset is returned in line 13.

#### Complexity analysis of REM-D

The theoretical complexity of REM-D (Algorithm 1) equals: number of classes × [(number of C5.0 calls × complexity of C5.0) + (number of hidden layers + 1) × [complexity of substitution + complexity of satisfiability checking + complexity of redundancy checking]]. The notation used throughout this analysis is as follows:

- *u*: umber of classes
- *k*: number of hidden layers
- *n*: number of data points
- *m*: number of features (i.e., neurons)
- *x*: maximum number of rules in each class ruleset
- *y*: maximum number of terms in each rule
- *z*: maximum number of substitutions for each term

Number of C5.0 calls (Algorithm 1): This number is equal to the sum of cardinality of set *T*, defined in line 6, across all layers. Set *T* consists of terms in the rules extracted between two adjacent layers proceeding the current layer *j* (line 4). Since there is only a single term in the initial 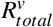, the cardinality of set *T* for the first hidden layer is 1. The C5.0 call for this term generate *n* − 1 new unique terms in the worst case scenario. Here is why: in the absence of pruning, the number of new terms is the same as the number of non-leaf nodes in the tree generated by C5.0. In order to calculate the maximum number of non-leaf nodes, we first look at the maximum number of leaf nodes. In the worst case scenario a tree induced between two layers has *n* leaves, where *n* is the number of data points. The total number of non-leaf nodes in a complete binary tree with *n* leaves is *n*− 1. Given that the number of new terms is the same as the number of non-leaf nodes of the tree, we end up with *n* − 1 terms in the worst case scenario. Since these newly generated terms will substitute the term in the initial 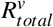, the number of unique terms in the newly formed 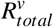 after substitution is equal to *n* − 1. This means that the cardinality of set *T* for layer *j* − 1 is *n* − 1 and hence C5.0 will be called *n* − 1 times. Each of the calls again will give rise to *n* − 1 new terms, the sum of which amount to (*n* − 1)(*n* − 1) = (*n* − 1)^2^. The continuation of this process makes the total number of C5.0 calls for a class equal to:

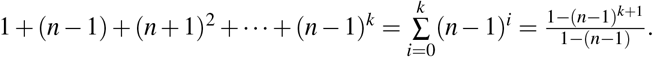

Complexity of C5.0 (Algorithm 1): C5.0 is a binary tree induction algorithm the complexity for which is known as *m* × *n*^2^ in the worst case scenario, where *n* is the number of data points and *m* is the number of features for each data point. In REM-D *m* refers to the number of neurons in layer *j* (line 4).

Complexity of substitution (Algorithm 2): complexity of this procedure depends on the total number of rules in a class (line 3) and the Cartesian product calculated in (line 8). Assuming that the maximum number of rules in each class ruleset, terms in each rule and substitutes for each term are *x, y* and *z*, respectively, the overall complexity of substitution procedure is *x* × *z*^*y*^.

Complexity of satisfiability and redundancy checking (Algorithm 1): We established *x* and *y* as the maximum number of rules in a ruleset and maximum number of terms within each rule, respectively. Satisfiability and redundancy checking both require iterating through every term in a rule for every rule, thus for each of this operations we have the complexity of *x*×*y*.

The overall complexity is thus: 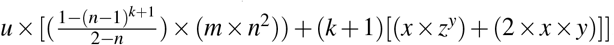. Determined by the dominant factor in *𝒪*(*u*[(*n*^*k*^*m*) + *k*[*xz*^*y*^ + *xy*]]) *n*^*k*^ and *z*^*y*^, the complexity grows exponentially with the number of layers and maximum number of terms in each rule. This is a limitation that can make rule extraction infeasible for large datasets. In practice however, C5.0 automatically does a considerable amount of pruning which makes the worst case scenarios avoidable. There are also interventions such as limiting the depth of trees between layers discussed earlier that can help with the empirical complexity. Other interventions include using parallel computing for multiclass problems to extract class rulesets simultaneously and using subsampling for large datasets, where only a fraction of *n* is used to build the trees between layers. In addition, feature sampling^55^ can be used to bound the number of *m* used for tree induction.

#### Comparison of REM-D with Zilke et al^39^

Zilke et al.^39^ uses C4.5 for tree induction and rule extraction between layers, while REM-D uses C4.5’s successor C5.0, due to the fact that C5.0 is more time and memory efficient and it extracts more accurate rulesets.

In^39^ first intermediate rules between all layers are extracted and stored, which requires a considerable amount of memory, and then they are subject to a final merge step. In favour of decreasing memory consumption, REM-D avoid storing all the intermediate rules. Instead, it merges the rules extracted between layers incrementally (e.g., rules extracted between layers 4 and 5 will be merged with rules extracted between layers 3 and 4 as soon as the latter become available).

Unlike^39^, REM-D does not extract rules from layers in isolation to one another. To improve time and memory consumption, after each step of merge, REM-D checks the newly formed set of rules for satisfiability and redundancy and only considers the non redundant terms within specifiable rules in the next rule extraction step. This essentially means dealing with less number of rules and also shorter ones.

### 4.4 REM-T rule extraction methodology

REM-T extracts rules (Algorithm 3) for a classification problem with *n* data points, 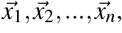, each with an associated class *y*_*i*_ ∈ {*y*_1_, *y*_2_, …, *y*_*u*_}. Rules are extracted from a trained random forest or decision tree. Here we explain the rule extraction from random forest, the procedure for decision tree is identical to that of random forest with a single tree.

The algorithm starts by iterating over each class (line 1). A total rule is introduced for each class (line 2). For each tree in the random forest (line 3), the branches of the tree (line 4) are traversed. If the branch ends at a leaf node that has the same label as the class (line 5), the algorithm creates a rule from it by the conjunction of conditions in each node in the branch and adds the created rule to the total ruleset for the class (line 6). Similar to Algorithm 1 (line 14) redundant terms are deleted (line 10). The total rulesets for each class are combined to give the overall ruleset (line 12) that is returned as output (line 13).

### 4.5 Rule scoring and ranking

The basis of personalised rule scoring and ranking is the score proposed by Mashayekhi et al.^27^,

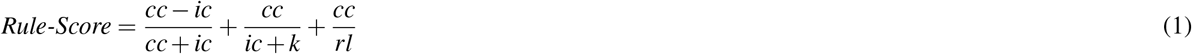

where *cc* and *ic* are the number of training samples covered by the rule and classified correctly and incorrectly, respectively, by the rule. *rl* denotes the rule length. *k* is set to 4 as per original work in Mashayekhi et al.^27^. However, other positive values can be used for *k* as its role is mostly to avoid the denominator becoming zero, and no significant change in the results is observed by modifying *k*.

#### Algorithm 3 REM-T

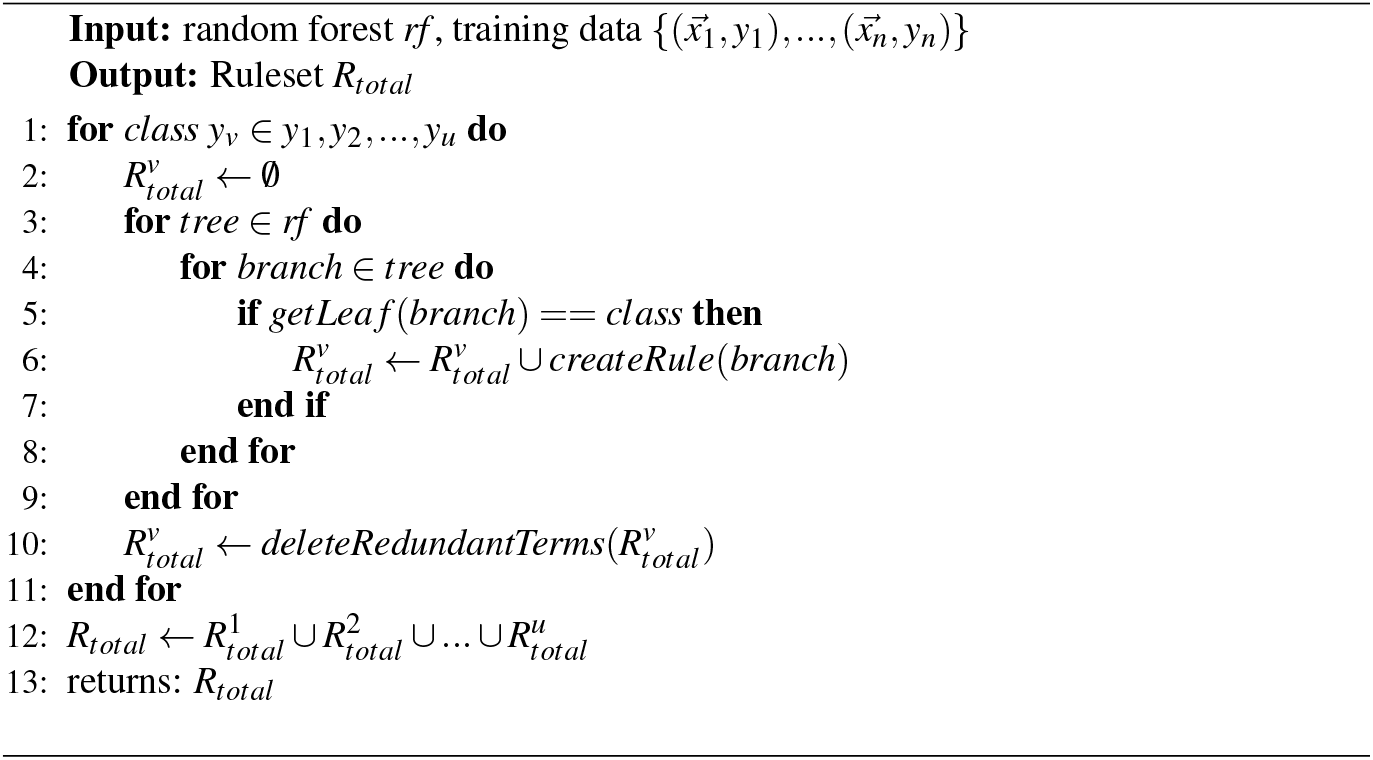

In the personalised rule scoring and ranking proposed by Müller^28^, where the assumption is that a list of preferred features is in hand, the equation is extended as follows:

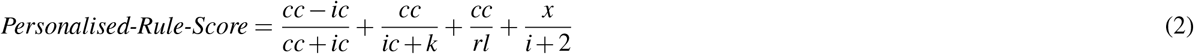

where *i* refers to the feature’s index in the list of preferred features: the lower the index, the more important is the feature. For a fixed *x*, the first feature (index 0) adds *x/*2 to the score, while the second feature (index 1) adds less (*x*/3) and so on. The positive constant *x* can be tuned to give the desired impact.

In the ranking imposed in case study II (Figure 4) we assumed all features are equally preferred (i.e., *i* = 0) and experimented with *x* ranging from 1 to 5 (Figure 6). Some values of *x* such as 4 and 5 give the most consistent results across all features in terms of increasing the number of rules that include the preferred features, while others are less consistent and may give more boost to only a certain feature. The results presented in Figure 4 use *x* = 5 in personalised ranking.

**Figure 6.**
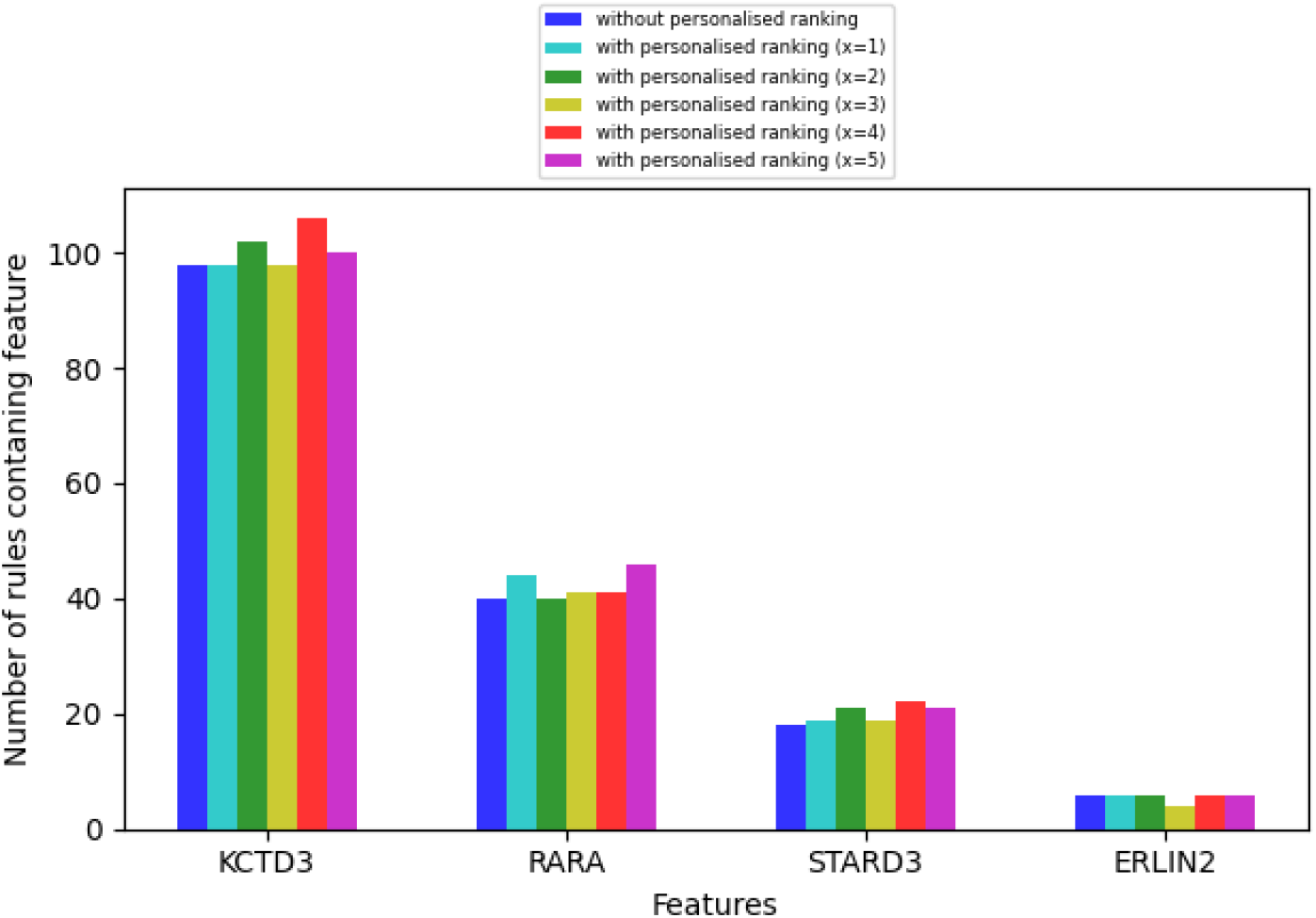
The impact of personalised ranking on increase in the number of higher rank rules (top 70% of the rules) that include the preferred features picked by clinicians, where *x* ranges from 1 to 5.

## Data and Code availability

All of the data and code necessary to reproduce our experimental findings cab be found at https://github.com/ZohrehShams/IntegrativeRuleExtractionMethodology.

## Acknowledgements

This work was supported by The Mark Foundation Institute for Integrated Cancer Medicine (MFICM). MFICM is hosted at the University of Cambridge, with funding from The Mark Foundation for Cancer Research (NY, U. S. A.) and the Cancer Research UK Cambridge Centre [C9685/A25177] (UK). We thank Dr. Oscar Rueda, Dr. Wei Cope and Mateo Espinosa Zarlenga for the helpful feedback and discussions on the work presented in this paper.

## Author Contributions

ZS initiated the study. ZS, NS, HT, PS, JA, PL, and MJ designed the study. ZS, BD, and SK designed the methods. ZS, SK and UM implemented the methods. ZS, BD, and PL performed analysis. All authors analyzed the results. ZS, BD, NS, MJ and PL wrote the manuscript. All authors reviewed and refined the manuscript.

## Supplementary materials

### Abbreviations and essential definitions

#### Abbreviations

ML: Machine Learning
MR: Machine Reasoning
DNN: Deep Neural Network
REM: Rule Extraction Methodology
REM-D: Rule Extraction Methodology from Deep Neural Networks
REM-T: Rule Extraction Methodology from Trees

## Essential Definitions

**Interpretability:** The degree to which “how” a model makes a prediction is understandable

**Explainability:** The degree to which “why” a model makes a prediction is understandable

**Bench-to-bedside:** the process of translating the results of research done in the laboratory into developing new ways of treating patients

**Care pathway:** A patient’s clinical plan of care

**Feature importance:** Set of domain features (i.e. piece of data or evidence) that contributed the most to a prediction

**Verifiability:** The degree to which the biological relevance of a model can be determined

**Simulatability:** The ease of generating a prediction for an input by following the operation of a model

**Reasoning:** Applying rules towards making predictions

**Accuracy:** The degree of alignment between the predictions of an extracted ruleset model and ground truth

**Fidelity:** The degree of alignment between the predictions of an extracted ruleset model and the predictions of the original model they are extracted from

**Efficiency:** Time and memory consumed to extract ruleset models

**Comprehensibility:** The number of rules in a ruleset and their average length

### Rule extraction details for case study I

Table 3 shows that the neural network model outperforms the decision tree model and the random forest model in terms of AUC, despite their high accuracy. The parameters used for the neural network model were outlined in the Method section. The random forest model used 50 trees each with maximum depth of 10 and half of the features considered at every split. The decision tree model also used a maximum depth of 10. Class weights were considered in both random forest and decision tree models.

Table 4 displays the details of rule extraction using REM-D for case study I reported in the Results and Method sections.

### Rule extraction details for case study II

Tables 5 and 6 show the details of rule extraction using REM-D and REM-T from mRNA and clinical data as outlined in the case study II, while Table 7 displays the details of their integration.

**Table 6.** Rule extraction from decision tree with 13 clinical variables as input and ER+/ER-as output.

| fold | REM-T<br>accuracy | size of<br>ruleset | rules<br>average length |
| --- | --- | --- | --- |
| 0 | 0.768 | 28 | 4.6 |
| 1 | 0.788 | 30 | 4.7 |
| 2 | 0.803 | 29 | 4.6 |
| 3 | 0.848 | 29 | 4.6 |
| 4 | 0.841 | 30 | 4.4 |
| average | **0.81** | **29.2** | **4.6** |

**Table 7.** Rule integration from neural network and decision tree.

| fold | REM-D + REM-T<br>accuracy | size of<br>ruleset | rules<br>average length |
| --- | --- | --- | --- |
| 0 | 0.927 | 206 | 5.0 |
| 1 | 0.952 | 133 | 5.0 |
| 2 | 0.952 | 211 | 5.3 |
| 3 | 0.922 | 43 | 4.0 |
| 4 | 0.911 | 47 | 3.9 |
| average | **0.933** | **128** | **4.7** |

### Interpretation of the results using enrichment analysis

Table 8 lists the genes that are occurring the most in the overall rulesets and rulesets for each class across five folds, when predicting histological types (Section 2.1 - Step d.1 of Figure 2). Genes listed for class IDC are passed on to g:Profiler^23^ for functional profiling using “Ordered query” option and “Data sources” set to “GO biological process” and “Reactome”. The detailed results are presented in Figure 7, which is an expansion of abstract results presented in Step d.2 of Figure 2. The connection between pathways is visualised using EnrichmentMap^24^ with the False-Discovery Rate (FDR) q-value cutoff set to 0.01. The visualisation is displayed in Figure 8, where similar pathways representing major biological themes are clustered automatically using “AutoAnnotate”^56^ on the basis of word frequency within the pathway names. As highlighted in Step d.3 of Figure 2, the main annotation found automatically are “Cycle phase cell” and “Telomere organization maintenance”. Figure 8 includes the name of pathways within each cluster, studying of which shows how annotations are extracted using the word frequency within the pathway names.

**Table 8.** Extension of the list in Figure 2d.1: Top 50 most recurring genes in rulesets across five folds.

|  |  |
| --- | --- |
| Top 50 features overall | **CDH1, AURKA, FOXM1, PSMD7, HYOU1, GRB7, EDC4, CSPP1, PPP1R14B, CIAPIN1, TOP2A, FAM91A1, UQCRB, NIP7, SAMM50, ROGDI, MCM10, NUBP1, ACTL6A, COG2, RCOR3, JOSD1, GTPBP4, TMEM219, COX4I1, GGPS1, DYNLL2, METRN, POP1, MRPS7, PARN, PPP2R5A, MAPK3, ALDH9A1, RFWD3, PEX11B, ZNF622, SLC35B1, OGFOD1, WDR68, TBCE, SRP9, ATP6V1H, CCNE2, BCL7C, STX3, TTPAL, HEATR3, DUS2L, BOP1** |
| Top 50 features for class IDC | **CDH1, AURKA, FOXM1, NIP7, HYOU1, CIAPIN1, PEX11B, SRP9, RFWD3, PARN, MCM10, UQCRB, HEATR3, GTPBP4, TMEM219, ALDH9A1, PPP2R5A, CCNE2, TTPAL, TOP2A, EIF3E, PSMD7, SAMM50, CYHR1, SNRPA1, RAD51C, KIAA0146, DUS2L, RFWD2, THOC6, TOM1L1, TLK2, GRB7, IMPAD1, EDC4, CCNE1, FAM91A1, FLYWCH2, GPR172A, NADSYN1, PSMC6, TERF1, JOSD1, RRS1, FAM96B, USF1, NSMCE2, DNAJA2, RAB25, BCKDK** |
| Top 50 features for class ILC | **CDH1, FOXM1, PSMD7, AURKA, GRB7, EDC4, PPP1R14B, CSPP1, HYOU1, FAM91A1, ROGDI, TOP2A, CIAPIN1, ACTL6A, NUBP1, COG2, RCOR3, UQCRB, SAMM50, COX4I1, GGPS1, JOSD1, DYNLL2, METRN, POP1, MRPS7, MAPK3, ZNF622, SLC35B1, OGFOD1, WDR68, TBCE, ATP6V1H, MCM10, STX3, BOP1, GTPBP4, BCL7C, MRPL28, TMEM219, PSMC2, MAF1, ANKRD11, PDSS1, NUDT5, PPP2R5A, LTV1, ALG1, GPR172A, DUS2L** |

**Figure 7.**
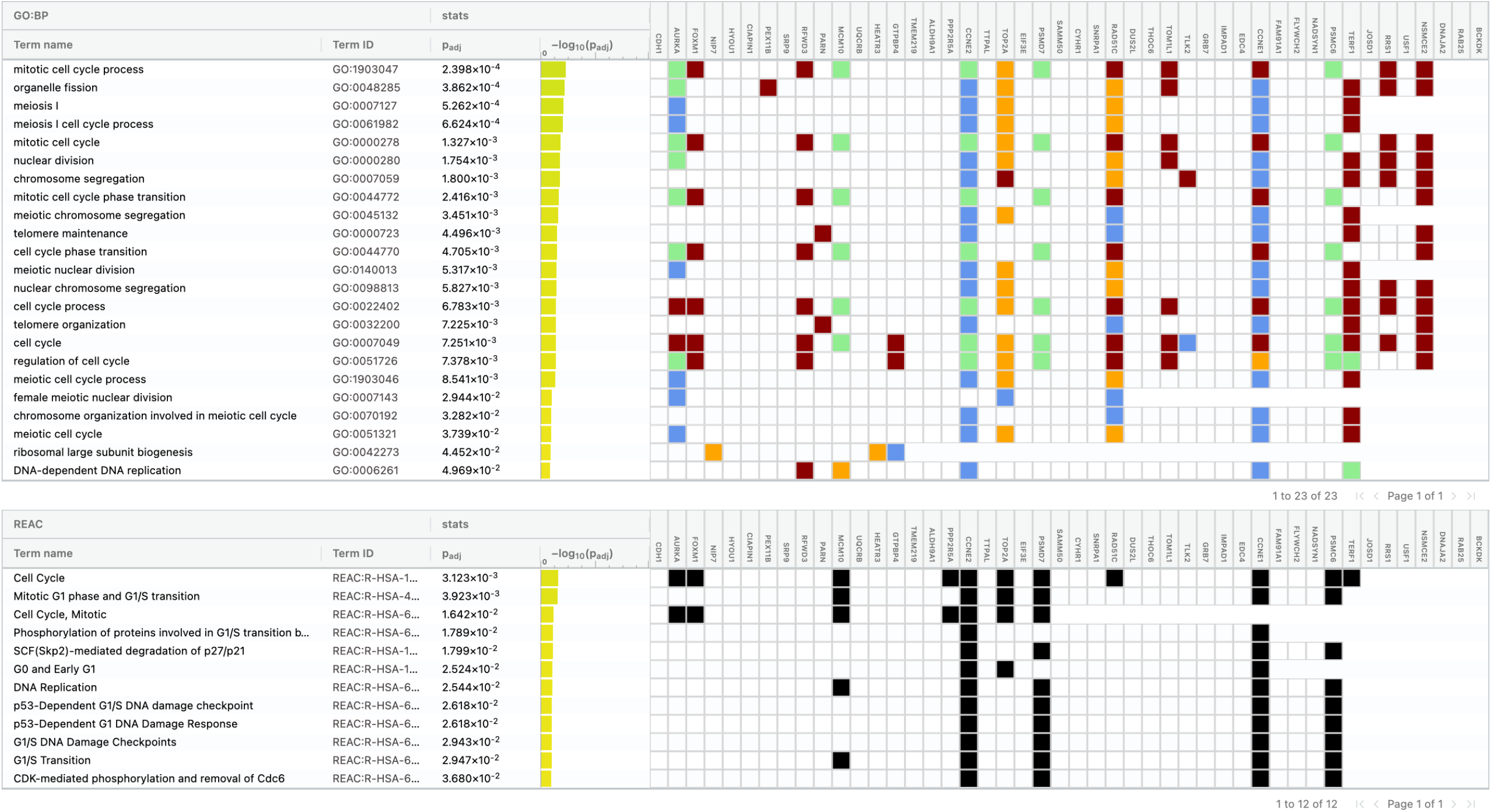
Functional profiling of top genes associated with class IDC using g:profiler. The terms listed under GO:BP correspond to enriched biological processes of GO displayed as orange circles in Figure 2d.1, while the terms under REAC corresponds to the enriched pathways of Reactome. Genes participated in the enrichment process of each term are highlighted accordingly in the right side of the figure.

**Figure 8.**
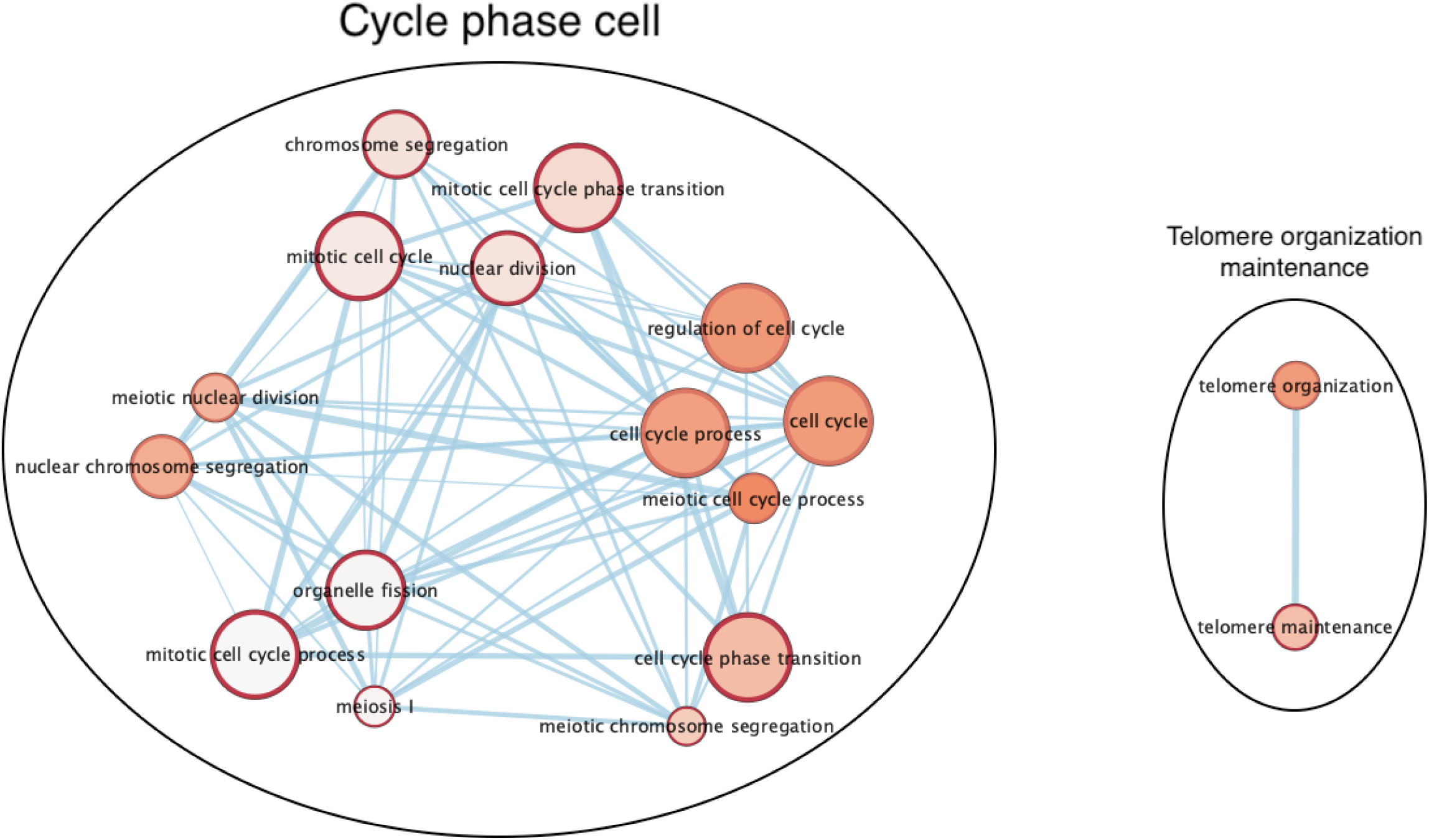
Visualisation of enrichment map based on functional profiling of top genes associated with class IDC, demonstrated in Figure 7. Clustering the overlapping pathways identifies two major biological themes, namely “Cycle phase cell” and “Telomere organization maintenance”.

### Footnotes

1 List of abbreviations and essential definitions can be found in the supplementary materials.

a From here onward, for succinctness we refer to all medical and clinicians as clinicians.

